# Lactylation-related genes prognostic model for triple-negative breast cancer and validation of a lactate–PFKP lactylation–glycolysis positive feedback loop that promotes malignant phenotypes

**DOI:** 10.64898/2026.11.20.768100

**Authors:** Saifeng Wang, Zicheng Zhang, Jiale Chen, Minfeng Liu

## Abstract

Triple-negative breast cancer (TNBC) is an aggressive breast cancer subtype associated with unfavorable clinical outcomes and a shortage of therapeutic targets. By integrating bioinformatic analysis of GEO datasets, we identified a four-gene lactylation-related signature (PFKP, VEGFA, PLOD2, and NDRG1) that effectively stratified TNBC patients by survival risk. In both the training and independent validation cohorts, the signature showed robust prognostic utility, with high-risk patients having worse overall survival. Among the signature genes, PFKP exhibited markedly increased lactylation in TNBC tissues. Sodium lactate treatment enhanced PFKP lactylation and enzymatic activity, whereas sodium oxamate reduced both. K688 was identified as the critical lactylation site on PFKP, and the K688R mutant showed decreased lactylation and enzyme activity. Functionally, PFKP lactylation promoted TNBC cell proliferation and migration while suppressing apoptosis, whereas the K688R mutant failed to restore these malignant phenotypes. These findings establish a lactylation-related prognostic signature for TNBC and reveal a lactate–PFKP lactylation–glycolysis feedback axis that may serve as a potential therapeutic target.

## 1. Introduction

Triple-negative breast cancer (TNBC) is a particularly aggressive breast cancer subtype, defined by the absence of estrogen receptor (ER), progesterone receptor (PR), and HER2 expression. It represents roughly 15–20% of breast cancer cases and is marked by rapid relapse, early metastasis, and poor survival^[1, 2]^. The absence of these key receptors limits the use of endocrine and anti-HER2 targeted therapies. Clinically, this leaves systemic cytotoxic chemotherapy as the primary treatment. However, responses are often short-lived, and chemoresistance remains a common clinical obstacle^[3]^. Although several prognostic models that rely on gene expression signatures have been constructed for TNBC, their predictive performance remains limited, and few have been translated into clinical practice. Therefore, there is a pressing demand to discover robust survival predictors and novel candidates for targeted therapy.

Metabolic reprogramming is a pivotal topic in oncology. A core characteristic of malignant cells is the Warburg effect, a process where glucose is converted into lactate irrespective of oxygen^[4]^. In addition to fulfilling the energy and synthesis needs of quickly reproducing cells, this metabolic change causes a significant buildup of lactate in the tumor microenvironment. Lactate, which was formerly thought to be a metabolic waste product, has emerged as an important signaling mediator that promotes aggressive tumor phenotypes, angiogenesis, and immune evasion^[5, 6]^. Recently, researchers identified lactylation as a newly described post-translational modification. This modification refers to the covalent attachment of a lactyl moiety to lysine residues within histone and non-histone proteins^[7]^. It is driven by intracellular lactate levels and serves as a key bridge linking cellular metabolism to epigenetic control. Lactylation of histones exerts direct control over gene expression^[8]^, while lactylation of non-histone proteins may modify their function, stability, and interactions with other proteins^[9, 10]^. Nevertheless, how lactylation contributes to TNBC biology and affects clinical outcome remains largely unexplored.

Among the potential targets of the lactate–lactylation regulatory axis, glycolytic enzymes represent particularly compelling candidates. Glycolytic enzymes catalyze the sequential reactions of glycolysis, and their activities are critical for determining the flux of glucose into downstream metabolic pathways^[11, 12]^. Emerging evidence indicates that glycolytic enzymes, including PKM2, undergo lactylation that regulates their activity, thereby forming feedback circuits which potentiate the glycolytic phenotype^[13]^. Consistent with this paradigm, lactylation-driven positive feedback circuits have been identified in several malignancies, such as the glycolysis–H3K18la– TTK/BUB1B loop in pancreatic ductal adenocarcinoma and the AKR1B10–lactate–H3K18la– LDHA circuit in hepatocellular carcinoma^[14, 15]^. Building on these findings, we hypothesized that lactylation of the key glycolytic enzyme constitutes a similar feedback mechanism in TNBC: lactate promotes lactylation of the enzyme, which enhances its activity, thereby accelerating glycolysis and further lactate accumulation, ultimately driving malignant progression.

Here, we show that lactylation of a critical glycolytic enzyme augments its catalytic activity, creating a self-reinforcing positive feedback circuit that promotes malignant phenotypes in TNBC. Our findings reveal a previously unrecognized lactylation-mediated mechanism contributing to TNBC aggressiveness and highlight a potential therapeutic axis.

## 2. Results

### 2.1. Identification of hub lactylation-related genes in TNBC

WGCNA was employed on the GSE21653 dataset to find the genes most closely linked to the prognosis of TNBC. The soft-thresholding power was set at 6 to yield a scale-free network (R^2^ ≥ 0.85) (Fig. 1A, B). Following module identification via the topological overlap matrix (TOM), the turquoise module was most strongly positively associated with TNBC status (correlation coefficient = 0.82, p = 1e^-24^), containing 1826 genes (Fig. 1C, D). The association of gene significance (GS) with module membership (MM) in the turquoise module (cor = 0.86, p < 1e^-200^) confirmed the validity of these genes (Fig. 1E).

**Figure 1.**
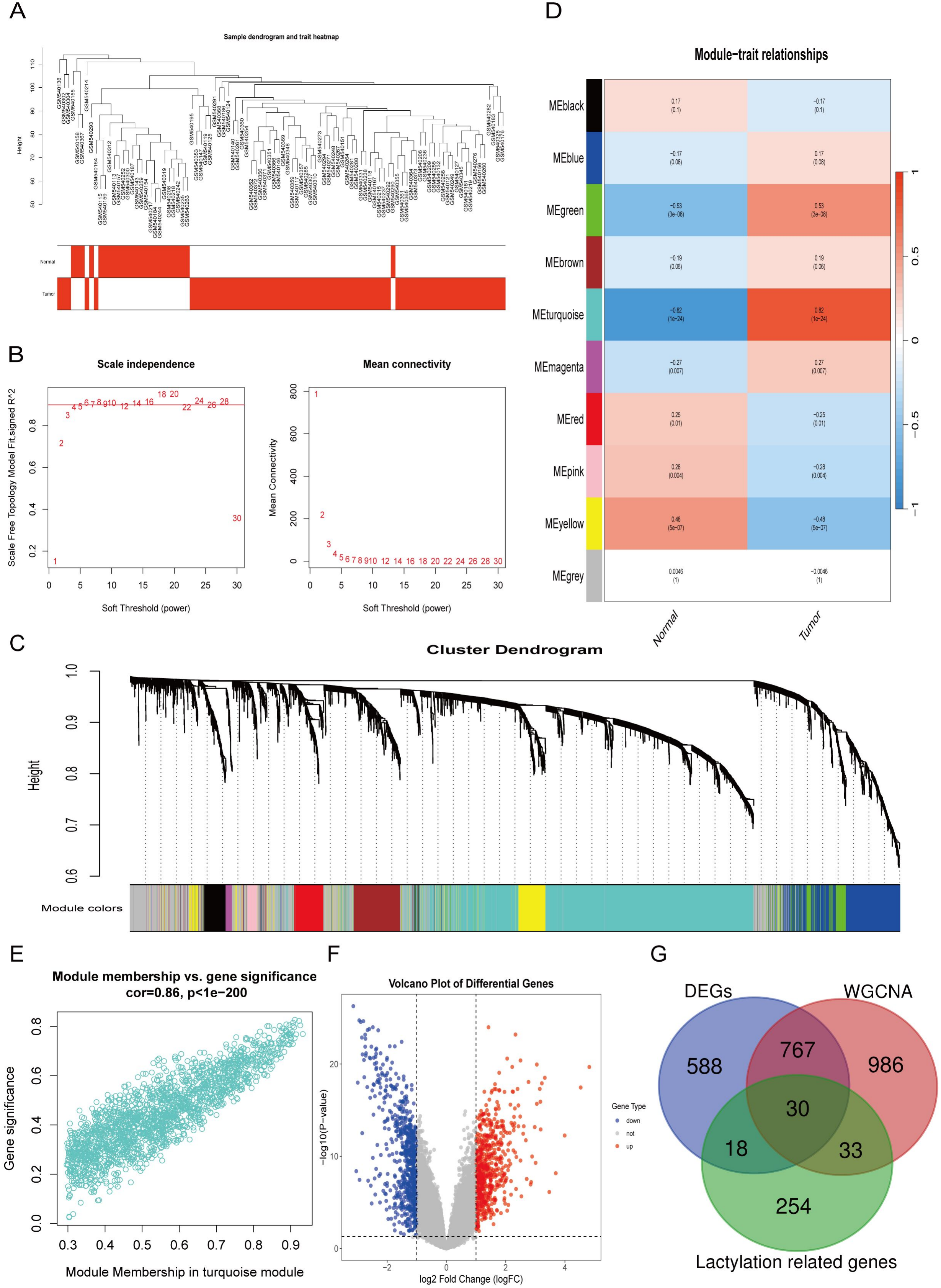
Identification of lactylation-related hub genes in TNBC via WGCNA and DEG analysis (A, B) Determining an appropriate soft-thresholding power for Weighted Gene Co-expression Network Analysis (WGCNA). (C) Dendrogram of gene clustering constructed via topological overlap matrix (TOM) dissimilarity. (D) Heatmap depicting associations between co-expression modules and clinical characteristics. (E) The connection between gene significance (GS) and module membership (MM) in the turquoise module. (F) Differentially expressed genes (DEGs) between TNBC and normal samples are shown in a volcano plot. (G) Venn diagram depicting the overlap among DEGs, lactylation-related genes (LRGs), and WGCNA turquoise module genes.

Based on the GSE21653 dataset, we compared TNBC samples with normal ones and identified 1403 differentially expressed genes (DEGs). Specifically, this comprised 661 upregulated and 742 downregulated genes with |log_2_(fold change)| > 1 and adjusted p < 0.05. The expression profiles are visualized in a volcano plot. (Fig. 1F). The intersection of these DEGs, 335 lactylation-related genes (LRGs), and the 1826 WGCNA turquoise module genes yielded 30 candidate hub lactylation-related genes (Fig. 1G).

### 2.2. Construction and validation of a prognostic lactylation-related gene signature

LASSO Cox regression was applied to the 30 candidate genes to determine which genes were most strongly linked to poor prognosis. Based on the optimal λ value and the minimal partial likelihood deviation, a four-gene model was chosen. (Fig. 2A, B). Risk scores were derived according to the formula below: Risk score = 0.0570 × exp (PFKP) + 0.1432 × exp (VEGFA) + 0.0759 × exp (PLOD2) + 0.0185 × exp (NDRG1).

**Figure 2.**
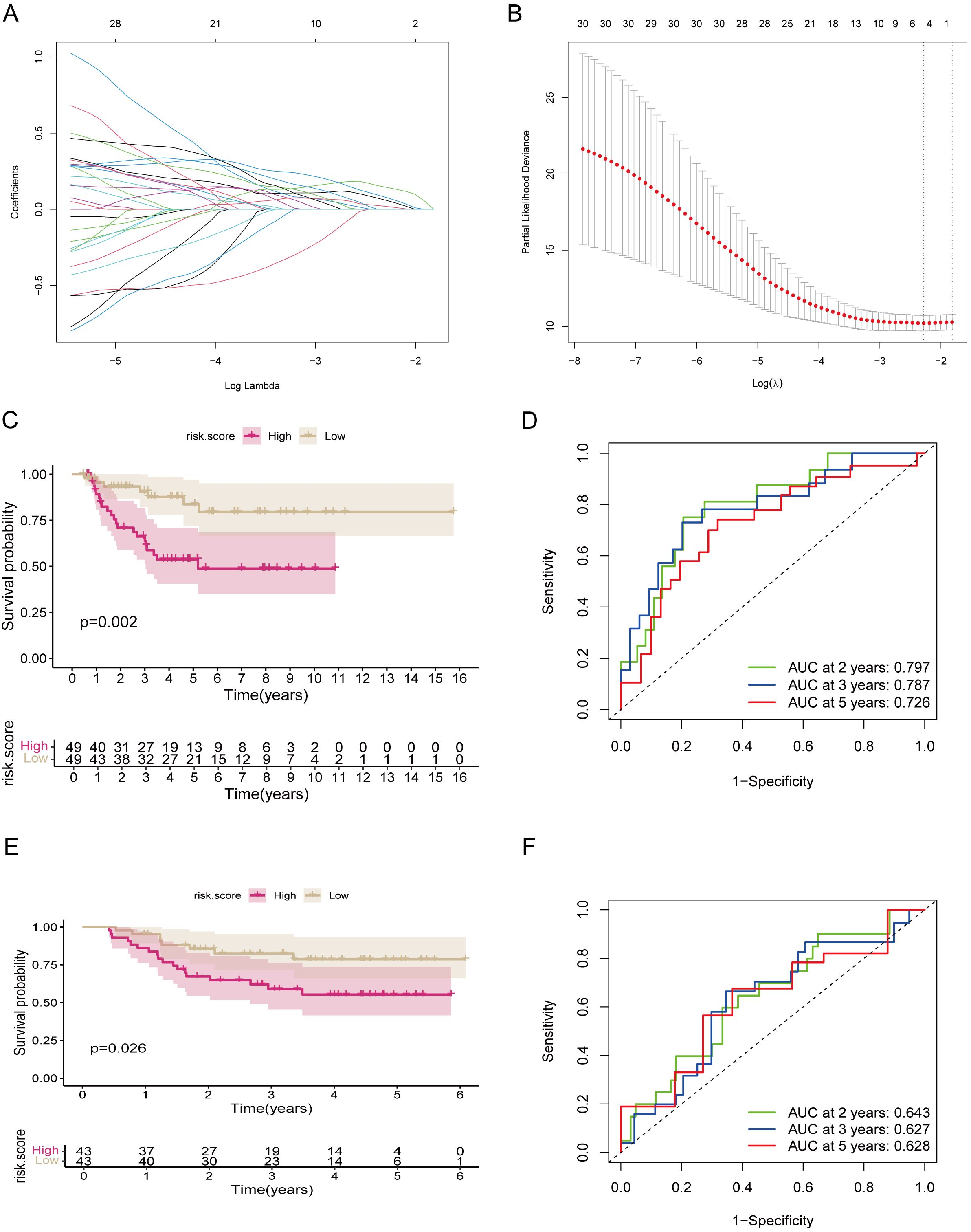
Construction and validation of a lactylation-related prognostic signature in TNBC. (A, B) LASSO Cox regression analysis performed on the GSE21653 training cohort. (C) Kaplan-Meier survival curves for high-risk and low-risk groups. (D) Time-dependent ROC curves assessing the 2-, 3-, and 5-year survival prediction accuracy. (E, F) External validation in the independent GSE25065 cohort: Kaplan-Meier survival curves (E) and time-dependent ROC curves (F).

Patients were classified into high-risk and low-risk cohorts using the median risk score as the threshold. Kaplan-Meier analysis showed that overall survival was substantially poorer in the high-risk group (Fig. 2C). The prognostic model showed satisfactory predictive accuracy, with time-dependent AUC values of 0.797, 0.787, and 0.726 for 2-, 3-, and 5-year survival, respectively (Fig. 2D). In the independent GSE25065 cohort, external validation supported the model’s robustness, with high-risk patients showing worse survival outcomes (Fig. 2E, F).

### 2.2. Immune cell infiltration landscape associated with the risk model

The lactate–lactylation axis acts as a bridge connecting metabolic reprogramming with immune regulation, remodeling the tumor immune microenvironment (TME). It involves driving macrophage polarization toward a pro-tumor (M2-like) phenotype and stabilizing regulatory T cells (Tregs)^[16]^. Lactylation also impairs dendritic cell antigen presentation and CD8^+^ T cell cytotoxicity while upregulating immune checkpoints^[17]^. Collectively, these effects create an immunosuppressive TME conducive to tumor immune escape and immunotherapy resistance. Given the known interplay involving lactate, lactylation, and the TME, CIBERSORT was employed to assess differences in immune cell infiltration across the risk groups (Fig. 3A, B). A pronounced enrichment of naïve CD4+ T cells, CD8+ T cells, naïve B cells, resting monocytes, resting dendritic cells, and resting mast cells was observed in the low-risk cohort. In contrast, the high-risk group showed higher proportions of resting macrophages (M0) and neutrophils (Fig. 3C-J). These results indicate that a distinct immunosuppressive landscape is a feature of the high-risk phenotype.

**Figure 3.**
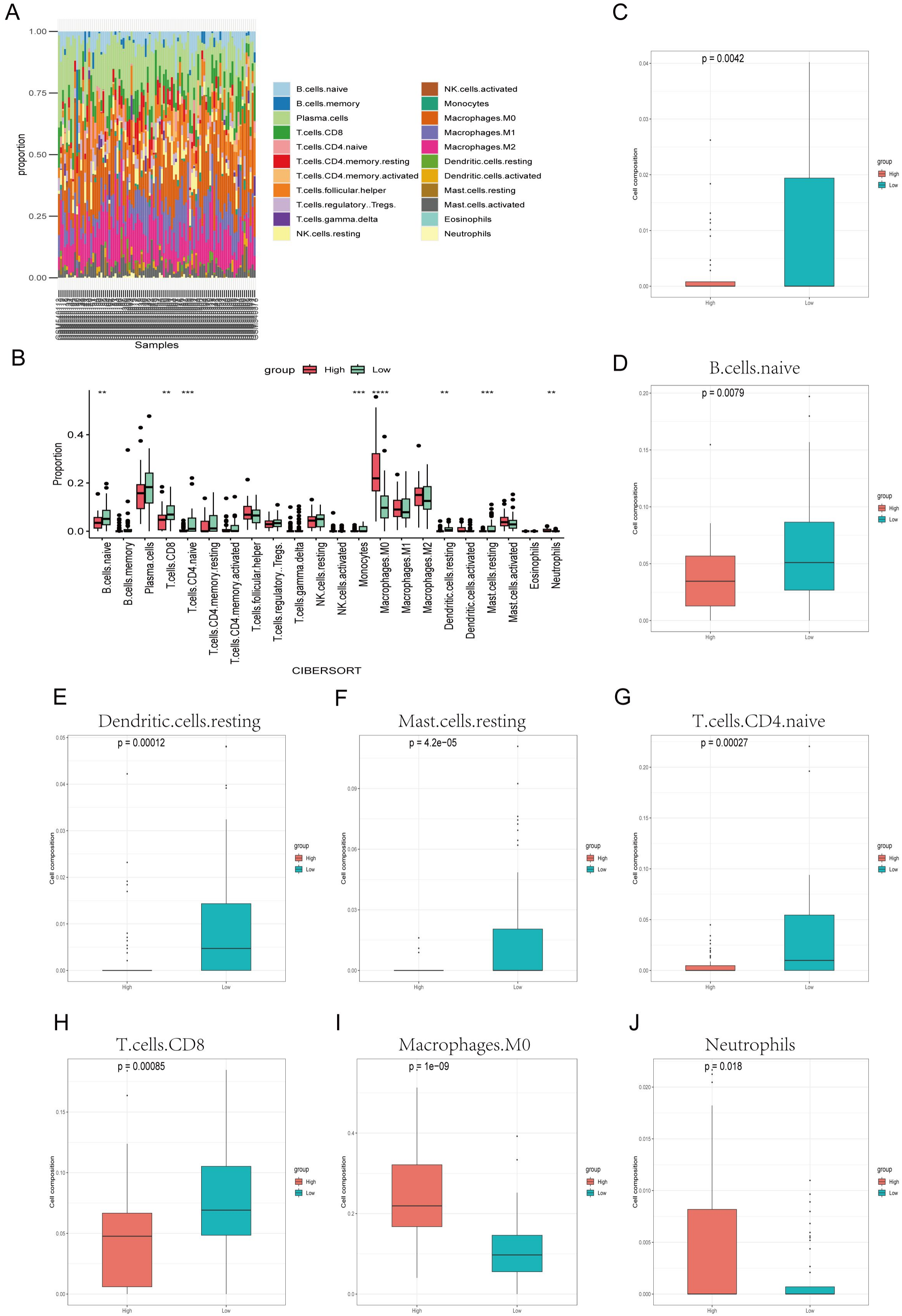
Immune cell infiltration landscape between high- and low-risk groups. (A, B) Immune cell composition profiles for the GSE21653 dataset, contrasting high-risk and low-risk groups using CIBERSORT. (C–J) Boxplots depicting the differential infiltration of specific immune subsets between the two risk groups.

### 2.4 Lactylation modification promotes malignant phenotypes in TNBC cells

To assess the functional impact of lactylation modification, we treated TNBC cells with sodium lactate to elevate lactylation or sodium oxamate to reduce it. Cell proliferative capacity and apoptosis were measured via CCK-8 and flow cytometry, respectively. The results revealed that high-level lactylation markedly boosted cell growth and suppressed apoptosis (Fig. 4A, C). Wound-healing assays further indicated that high lactylation conditions significantly increased cell migration (Fig. 4B). Collectively, these observations highlight a pro-tumorigenic role for lactylation in TNBC progression.

**Figure 4.**
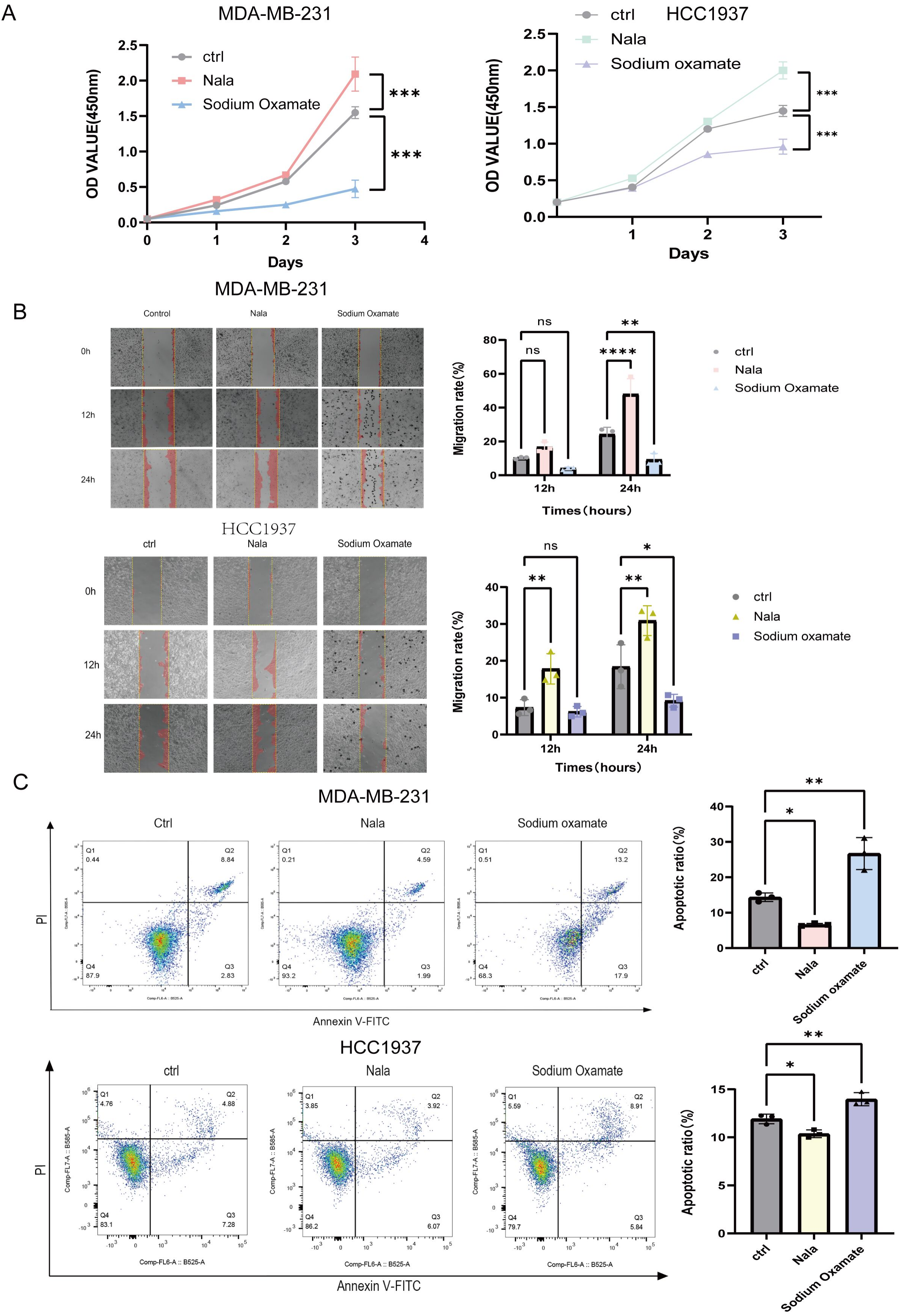
Lactylation modification promotes malignant phenotypes in TNBC cells. (A)CCK-8 assay showing proliferation of triple-negative breast cancer (TNBC) cells treated with sodium lactate or sodium oxamate to modulate lactylation. (B) Wound-healing assay showing migration of TNBC cells. (C) Flow cytometry analysis of apoptosis after Annexin V-FITC/PI staining. Values are expressed as the mean ± SD from three independent experiments. *p < 0.05,**p < 0.01, ***p < 0.001, and ****p < 0.0001 were considered statistically significant.

### 2.5. PFKP is a direct target of lactylation and its activity is modulated by lactylation

PFKP mediates the irreversible phosphorylation of fructose-6-phosphate (F6P) to fructose-1,6-bisphosphate (F-1,6-BP) during glycolysis, a critical point that determines the flux of glucose into downstream metabolic pathways^[18, 19]^. Increased expression of this protein has been correlated with unfavorable prognosis across multiple tumors^[20]^, including breast cancer^[21]^, suggesting a central role in supporting the anabolic and energetic demands of malignant cells. However, the exact molecular basis underlying PFKP activity in TNBC remains limited. Recent studies have demonstrated that lactate levels modulate protein expression or activity via lactylation, particularly affecting the core rate-limiting enzymes in glycolysis. We therefore hypothesized that PFKP is subject to regulation by lactylation. To address this, we collected tumor tissues from four TNBC patients, along with paired normal adjacent tissues (NAT), to determine the differences in protein lactylation levels. Mass spectrometry data indicated that PFKP lactylation was elevated by approximately 25-fold in tumor tissues compared to NAT. PFKP ranked among the top 10 proteins with the greatest increase in lactylation (Fig. 5A), while the other three model genes showed no significant changes. Therefore, we focused on PFKP lactylation.

**Figure 5.**
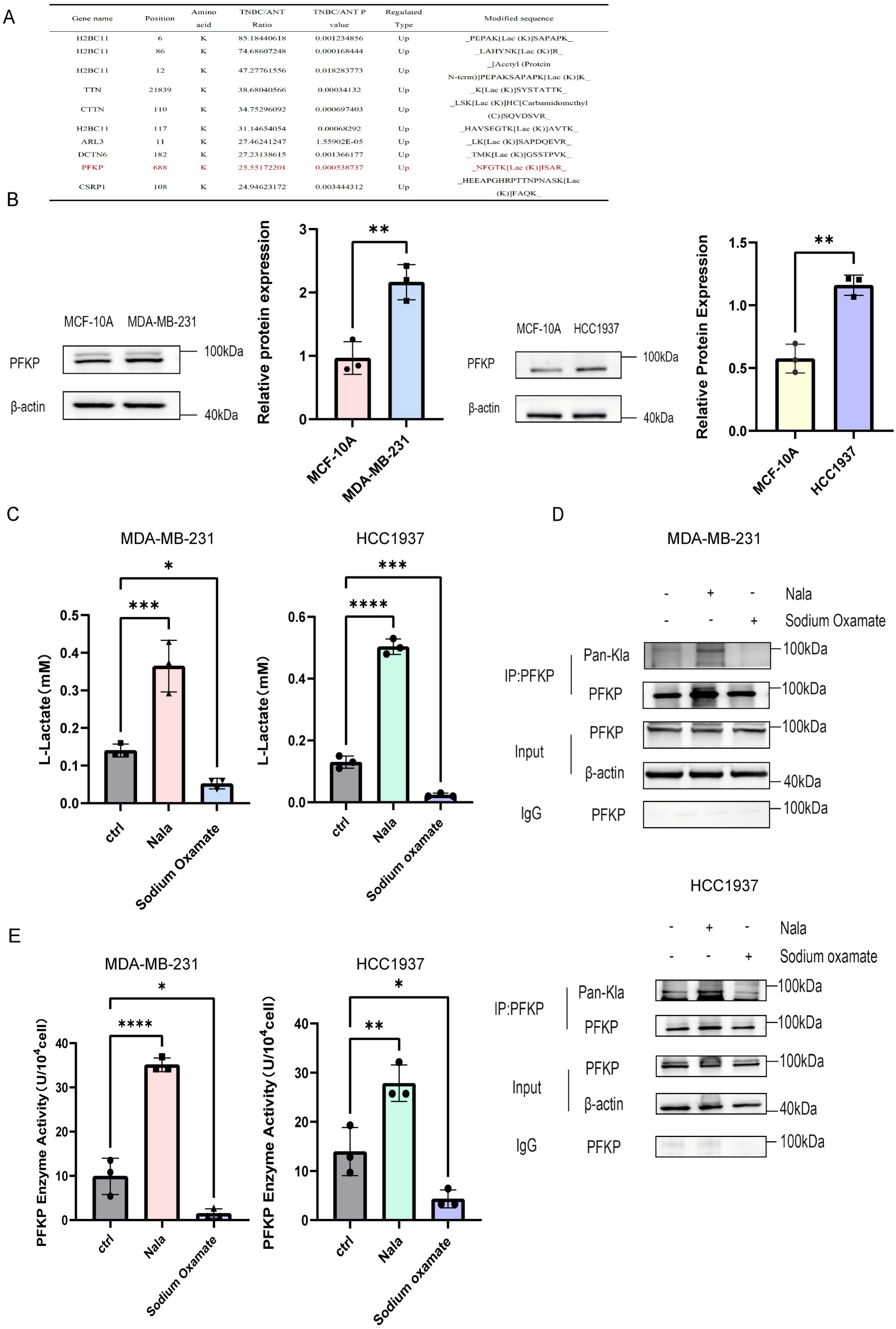
Identification of PFKP as a lactylation target in TNBC. (A) The top 10 lactylation sites exhibiting the most prominent increase in triple-negative breast cancer (TNBC) compared with normal adjacent tissues (NAT). (B) Western blot analysis displaying PFKP protein levels in TNBC cells versus normal MCF-10A cells. (C) Intracellular lactate levels in TNBC cells treated with 20 mM sodium lactate or 10 mM sodium oxamate for 24 h. (D) Co-IP of PFKP lactylation in TNBC cells treated with sodium lactate or sodium oxamate. (E) PFKP enzymatic activity in TNBC cells treated with sodium lactate or sodium oxamate. Values are expressed as the mean ± SD from three independent experiments. *p < 0.05, **p < 0.01, ***p < 0.001, and ****p < 0.0001 were considered statistically significant.

Aligning with the training cohort analysis, PFKP protein expression was observed to be markedly elevated in TNBC cells compared with MCF-10A cells (Fig. 5B). Treatment with sodium lactate (an exogenous lactate donor) significantly increased intracellular lactate level, while sodium oxamate (an LDHA inhibitor) decreased it (Fig. 5C). Co-IP experiments demonstrated that sodium lactate treatment upregulated PFKP lactylation, whereas sodium oxamate downregulated it (Fig. 5D). Importantly, PFKP enzymatic activity was enhanced by increased lactylation and suppressed by reduced lactylation (Fig. 5E). These results indicate that lactate levels directly influence both the lactylation modification and the enzymatic activity of PFKP.

### 2.6. K688 is the critical lactylation site on PFKP

Mass spectrometry identified lysine 688 (K688) as a lactylation site on PFKP. To validate this, a K688R mutant (lysine-to-arginine) that abrogates lactylation at this site was designed. MDA-MB-231 and HCC1937 cells were transfected with wild-type PFKP, K688R mutant, or empty vector (Fig. 6A). Co-IP confirmed that the wild-type PFKP exhibited robust lactylation, whereas the K688R mutant showed significantly reduced lactylation (Fig. 6B). The enzymatic activity of the K688R mutant was also markedly impaired compared to wild-type PFKP (Fig. 6C). Functional assays demonstrated that while wild-type PFKP overexpression enhanced cell viability, migration, and resistance to apoptosis, the K688R mutant failed to promote these malignant phenotypes (Fig. 7A-C). These results confirm that lactylation at K688 is essential for PFKP-mediated oncogenic functions in TNBC.

**Figure 6.**
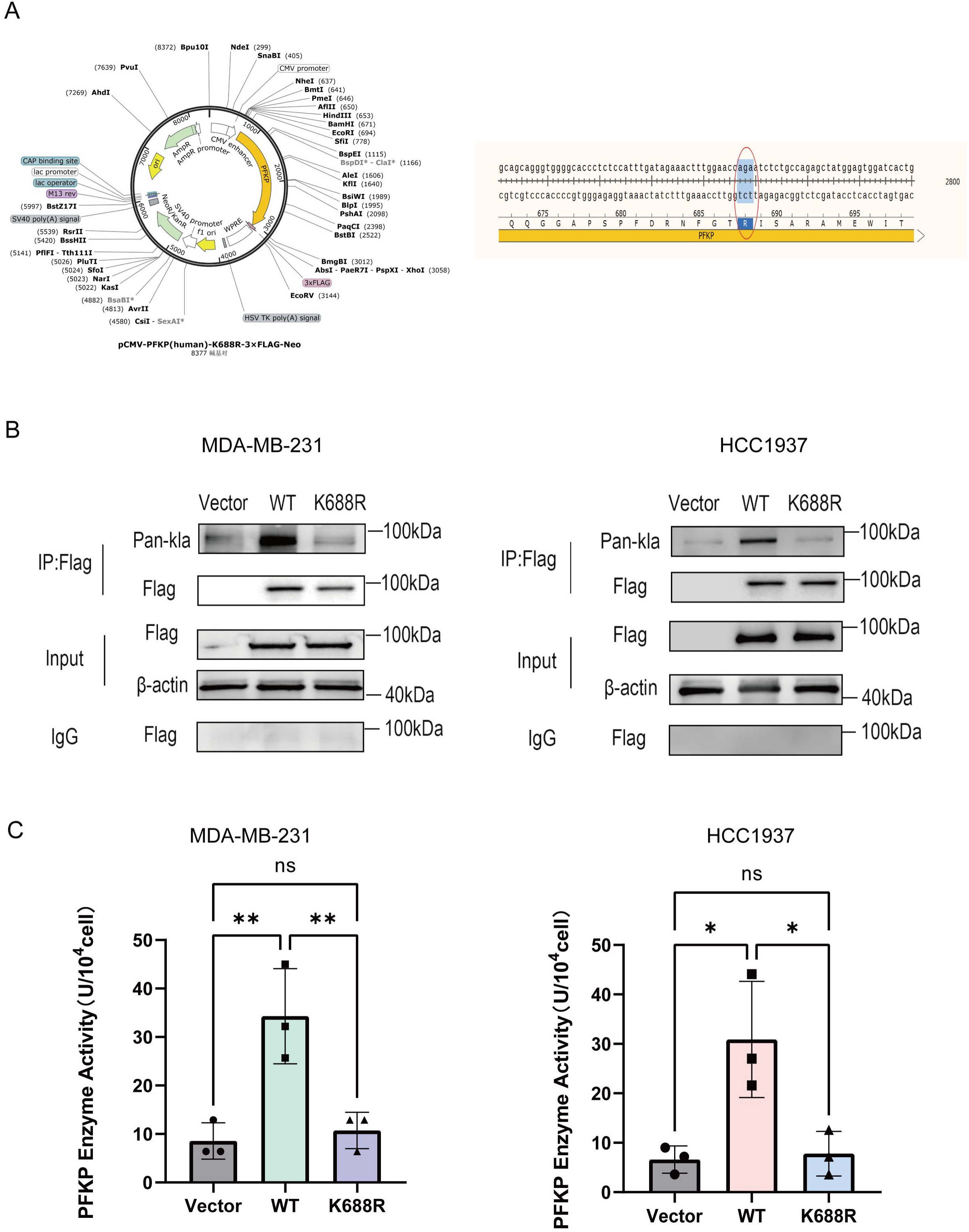
K688 is the critical lactylation site regulating PFKP enzymatic activity. (A) Schematic illustration of the plasmid structure of the K688R mutant, in which lysine at position 688 is replaced by arginine. (B) Co-IP analysis of lactylation levels in wild-type PFKP and the K688R mutant. (C) Enzymatic activity of wild-type PFKP and the K688R mutant. Values are expressed as the mean ± SD from three independent experiments. *p < 0.05, **p < 0.01, ***p < 0.001, and ****p < 0.0001 were considered statistically significant.

**Figure 7.**
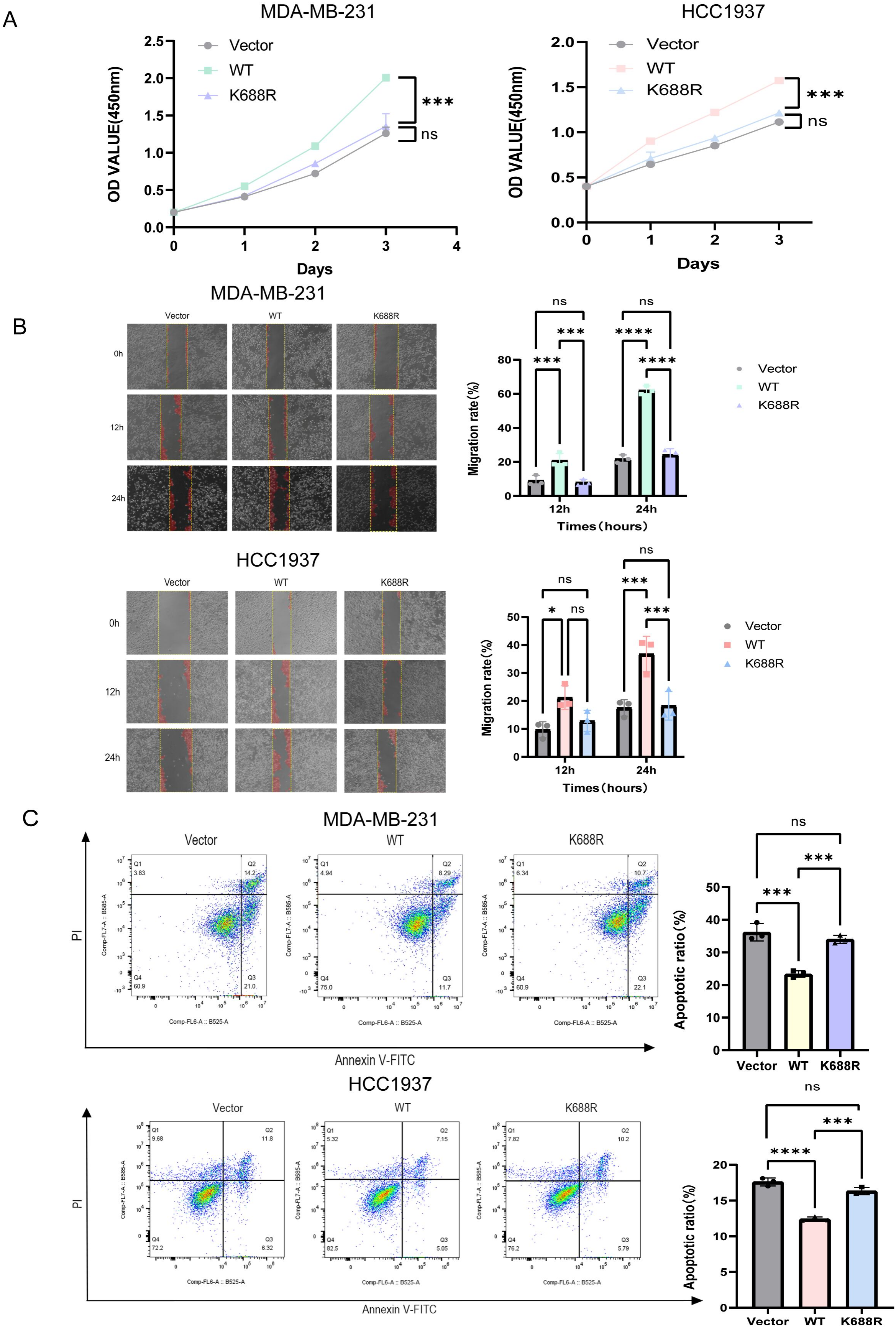
K688 lactylation is required for PFKP-mediated oncogenic phenotypes in TNBC. (A) CCK-8 assay showing cell viability of triple-negative breast cancer (TNBC) cells transfected with wild-type PFKP, K688R mutant, or empty vector. (B) Wound-healing assay showing cell migration. (C) Flow cytometry analysis of apoptosis. Values are expressed as the mean ± SD from three independent experiments. *p < 0.05, **p < 0.01, ***p < 0.001, and ****p < 0.0001 were considered statistically significant.

## 3. Discussion

In the present study, we developed a novel prognostic model based on a four-gene lactylation-related signature (PFKP, VEGFA, PLOD2, and NDRG1) that effectively stratifies TNBC patients by survival risk. More importantly, we elucidated a previously unrecognized mechanism by which the glycolytic enzyme PFKP is post-translationally modified by lactylation, a process that enhances its activity and promotes malignant progression of TNBC. Our findings collectively establish lactylation as a significant molecular determinant of TNBC prognosis and aggressiveness, offering new avenues for biomarker development and therapeutic targeting.

The signature developed in this study exhibits robust prognostic capability for overall survival in TNBC. Through LASSO Cox regression, we identified a four-gene signature that could accurately stratify patients into high-risk and low-risk groups. External validation in an independent cohort indicated its generalizability. The four genes comprising the signature (PFKP, VEGFA, PLOD2, and NDRG1) are all linked to hypoxia, glycolysis, and the tumor microenvironment, consistent with the functional consequences of lactate accumulation. This aligns with previous studies that have established LRG-based models for other malignancies, such as hepatocellular carcinoma and pancreatic cancer^[22, 23]^, reinforcing the idea that lactylation patterns can serve as a molecular signature for cancer prognosis. The elevated expression of these genes in TNBC samples compared to normal tissues further underscores their potential biological relevance in this aggressive subtype. PFKP governs the initial rate-limiting reaction of glycolysis, and its enzymatic activity is critical for directing glucose flux into energy production and biosynthetic pathways^[24]^. VEGFA, a key driver of angiogenesis, is upregulated in TNBC and is partially regulated by lactate and hypoxia^[25, 26]^. Similarly, PLOD2, a collagen-modifying enzyme, is frequently overexpressed in solid tumors and its expression is promoted by hypoxia, facilitating extracellular matrix remodeling and metastasis^[27, 28]^. NDRG1, a protein involved in cell differentiation and stress responses, is associated with tumor progression and adverse prognosis across multiple cancer types^[29]^. Collectively, the genes in our signature highlight the interplay among metabolic reprogramming, extracellular matrix remodeling, and angiogenesis, all of which are hallmarks of aggressive TNBC.

Functional experiments firmly established the oncogenic role of lactylation in TNBC. Treatment with sodium lactate to elevate lactylation markedly increased cell proliferation and migration while reducing apoptosis, whereas sodium oxamate-mediated reduction of lactylation produced the opposite effects. These observations support a tumor-promoting function of lactylation in TNBC progression and led us to seek the specific protein targets responsible for these effects. Through mass spectrometry analysis of paired TNBC and normal adjacent tissues, we identified PFKP as a highly lactylated protein in tumor tissues, ranking among the top 10 proteins with the greatest increase in lactylation. We further confirmed that lactate levels directly influence both the lactylation and the enzymatic activity of PFKP: sodium lactate treatment upregulated PFKP lactylation and enhanced its enzymatic activity, while sodium oxamate downregulated both. These results establish PFKP as a lactate-responsive lactylation target in TNBC. We then demonstrated that PFKP undergoes lactylation at lysine 688 (K688), a modification that enhances its enzymatic activity. The K688R mutant exhibited significantly reduced lactylation and impaired enzymatic activity compared with wild-type PFKP, and failed to restore the malignant phenotypes observed upon wild-type PFKP overexpression. These results confirm that K688 is the critical residue mediating PFKP lactylation and its oncogenic functions in TNBC.

This is a crucial finding, as it uncovers a self-reinforcing cycle: elevated lactate, a product of glycolysis, enhances PFKP lactylation, which in turn potentiates glycolysis and promotes lactate production, and then the accumulated lactate further drives PFKP lactylation. This mechanism is consistent with recent discoveries on other glycolytic enzymes. For example, PKM2 lactylation is known to facilitate tumor progression and suppress inflammatory responses in macrophages^[13]^. Similarly, Cheng et al. reported that ENO1 lactylation boosted its stability and enzymatic activity, thereby promoting glycolysis and lactate production in TNBC. Notably, this accumulated lactate further drives ENO1 lactylation, creating a positive feedback loop that fuels malignant progression^[30]^. The identification of a lactate-PFKP lactylation-glycolysis circuit may explain, in part, the highly aggressive nature of TNBC, as it demonstrates a self-sustaining mechanism that confers a survival advantage to tumor cells. Abolishing this modification via a K688R point mutation in PFKP impaired these oncogenic phenotypes. Our findings directly link the lactylation of PFKP to the aggressive traits of TNBC cells. Thus, PFKP serves not merely as a glycolytic enzyme but also as a pivotal component of a signaling pathway linking metabolic state to oncogenic signals.

Admittedly, our research has certain limitations. First, our prognostic model was constructed and verified utilizing publicly available microarray datasets. While this demonstrates its robustness, future studies using larger, more diverse, and prospectively collected cohorts are necessary to confirm its clinical utility. Second, our functional experiments were performed in a limited number of TNBC cell lines in *vitro*. However, the role of PFKP lactylation in tumor progression needs to be confirmed in *vivo*, such as in orthotopic xenografts, to better mimic the complexities of the tumor microenvironment. Third, how exactly K688 lactylation enhances PFKP enzymatic activity requires further investigation. Studies are needed to determine whether this modification alters PFKP’s affinity for its allosteric regulators or promotes its tetramerization. Finally, the enzymes responsible for PFKP lactylation should be explored to complete the mechanistic framework of this study and fully realize its therapeutic potential.

## 4. Conclusions

In summary, this study provides a dual contribution to the field of TNBC research. We have established a validated lactylation-related prognostic gene signature that can effectively identify TNBC patients at high risk of mortality. Furthermore, PFKP was identified as a functional lactylation target, revealing a targetable mechanism driving TNBC aggressiveness. K688 lactylation of PFKP enhances its enzymatic activity, perpetuating a glycolysis–lactylation positive feedback loop that promotes malignant phenotypes. These findings emphasize the importance of metabolic reprogramming and post-translational modifications in TNBC pathogenesis and could provide a foundation for new therapeutic strategies designed to target the lactate–PFKP lactylation–glycolysis axis. A schematic overview summarizing these core findings is presented in the Graphical Abstract (Fig. 8).

**Figure 8.**
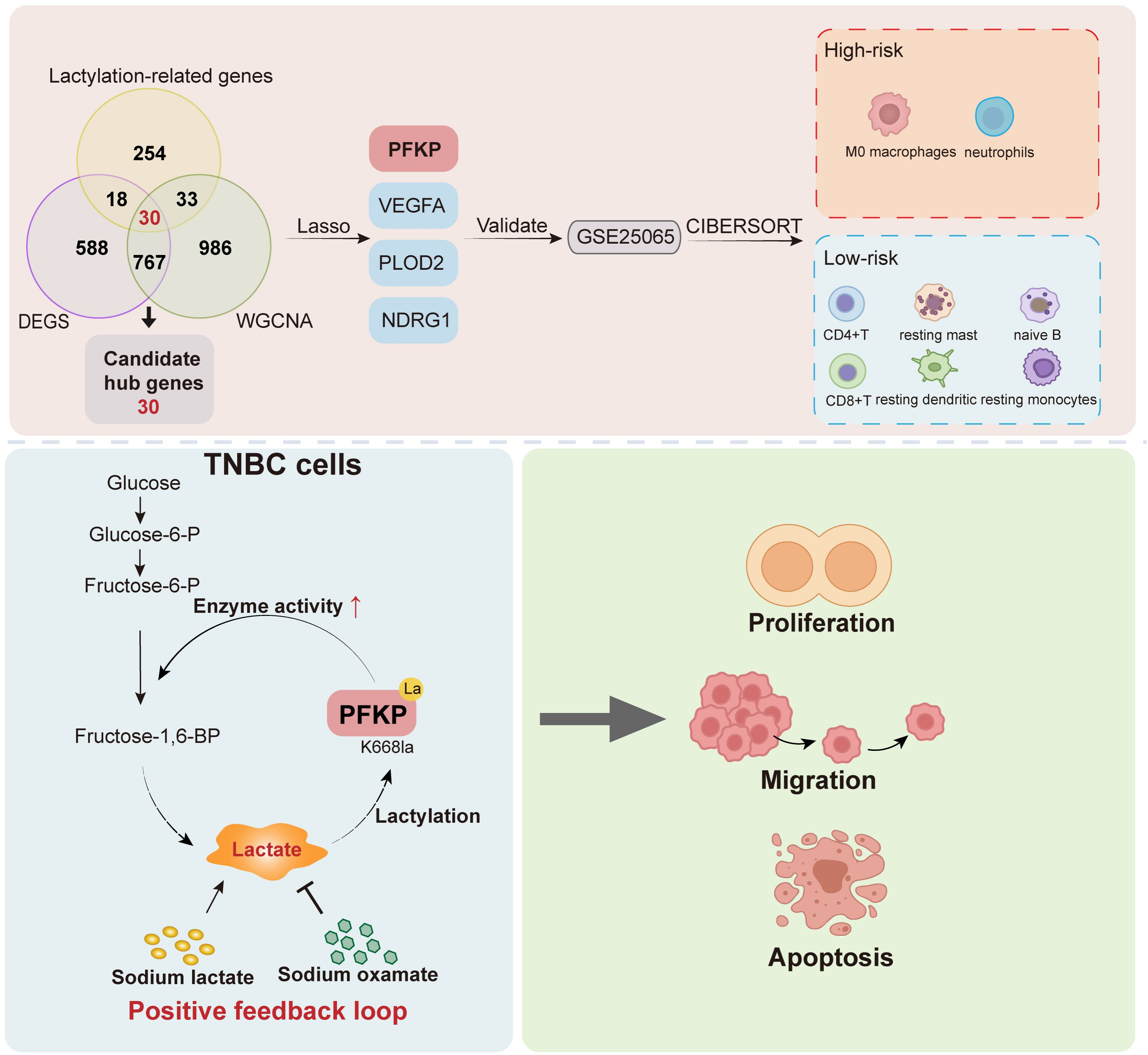
The Graphical Abstract.

## 5. Materials and Methods

### 5.1. Data Acquisition and Processing

We retrieved transcriptomic data and matched clinical records for TNBC patients from the Gene Expression Omnibus (GEO) database (http://www.ncbi.nlm.nih.gov/geo/). The GSE21653 dataset, consisting of 26 normal and 74 TNBC samples, was utilized as the training cohort, and GSE25065, comprising 19 normal and 67 TNBC samples, served as the independent validation cohort. A set of 335 lactylation-related genes (LRGs) (Appendix Table S1) was retrieved from the MSigDB^[31]^ (https://www.gsea-msigdb.org/gsea/msigdb/).

### 5.2. Identification of Differentially Expressed Genes (DEGs)

Differential expression analysis between TNBC and normal samples was performed using the limma package with criteria of |log_2_(fold change)| > 1 and adjusted p < 0.05^[32]^. The ggplot2 package in R was employed to create volcano plots displaying the DEGs

### 5.3. Weighted Gene Co-expression Network Analysis (WGCNA)

With the aim of detecting gene modules of clinical relevance, we performed weighted gene co-expression network analysis (WGCNA) via the corresponding R package^[33]^. The top 25% of genes with the highest median absolute deviation (MAD) from the normalized expression matrix (GSE21653, n = 110) were selected as input. A soft-thresholding power of 6 was selected so that the resulting network met the scale-free criterion (R^2^ ≥ 0.85). We characterized modules as gene clusters comprising a minimum of 30 genes. Each module was then represented by its eigengene, and we selected the one most strongly correlated with TNBC clinical traits.

### 5.4. Construction and Validation of the Prognostic Signature

We extracted the overlapping genes among DEGs, LRGs, and WGCNA-derived modules. Genes selected by univariate Cox regression were further analyzed by LASSO regression implemented in the glmnet R package (family = “cox”). To determine the optimal penalty parameter (λ), we applied the cv.glmnet function with 10-fold cross-validation (nfolds=10). The final model was established at lambda.min, which corresponded to the λ value yielding the minimal cross-validation error. Subsequently, a multivariate Cox regression model was fitted using LASSO-derived risk scores together with clinical variables. We calculated the risk score for each patient using the following formula: risk score = ∑_i_ Coefficient(gene_i_) × mRNA Expression(gene_i_). Patients were stratified into high-risk and low-risk groups according to the median risk score threshold. Kaplan-Meier survival curves and time-dependent receiver operating characteristic (ROC) curves were plotted utilizing the survival and timeROC packages, respectively. Finally, external validation of the prognostic signature was conducted in the GSE25065 dataset.

### 5.5. Immune Cell Infiltration Analysis

We utilized the CIBERSORT algorithm to quantify the relative abundance of immune cell subsets across samples from the GSE21653 dataset^[34]^. Furthermore, disparities in immune cell infiltration between the high-risk and low-risk cohorts were assessed via the Wilcoxon rank-sum test.

### 5.6. Lactic Acid Detection

The assay was performed using a lactate assay kit (S0208S, Beyotime) following the manufacturer’s protocol. Cells were lysed in extraction buffer, and following centrifugation of the resulting lysates, the supernatant was collected. A series of standard dilutions was prepared to establish a standard curve. Equal volumes of sample supernatants and standards were loaded into the reaction wells, the detection reagent was applied to every well. After mixing, the plate was incubated at 37°C for 30 minutes. Absorbance was measured at 450 nm on a microplate reader. For each sample, lactate concentration was derived from a standard curve constructed with the serial standards.

### 5.7. Cell Culture and Treatment

Human TNBC cell lines MDA-MB-231 and HCC1937 were cultured in Dulbecco’s Modified Eagle’s Medium (DMEM) supplemented with 10% fetal bovine serum (FBS) and 1% penicillin-streptomycin, while the normal breast epithelial cell line MCF-10A was cultured in MCF-10A Specialized Medium (CM-0525, Procell). All cells were maintained at 37°C in an atmosphere containing 5% CO2. For lactate modulation, cells were treated with 20 mM sodium lactate (S817836, Macklin) for 24 hours to elevate intracellular lactate, or with 10 mM sodium oxamate (KCM14317, KKL Med), a lactate dehydrogenase A (LDHA) inhibitor, to reduce lactate production. MDA-MB-231 and HCC1937 cells were transiently transfected using Lipofectamine 3000 with wild-type PFKP, K688R mutant PFKP, or empty-vector plasmids (synthesized by MiaoLingBio). The K688R mutant was generated by site-directed mutagenesis (AAG→AGG), and the full sequences are provided in Dataset EV1.

### 5.8. Western Blotting and Co-Immunoprecipitation (Co-IP)

For extraction of total protein, RIPA buffer supplemented with protease and phosphatase inhibitors was used to lyse the cells. Co-IP was then carried out using the Protein A+G Magnetic Beads Kit (P2179S, Beyotime) strictly according to the instructions. Following overnight incubation at 4°C, the lysates were treated with anti-IgG, anti-FLAG (MA1-91878, Invitrogen), or anti-PFKP (13389-1-AP, Proteintech). Next, after boiling for 5 min at 95°C in SDS-PAGE loading buffer, bound proteins were eluted and then subjected to Western blotting. The samples were loaded onto 10% SDS-PAGE and transferred to PVDF membranes. After blocking with 5% non-fat milk, membranes were subjected to overnight incubation with primary antibody: anti-β-actin (K200058M, Solarbio, 1:10000), anti-Kla (PTM-1401RM, PTM Bio, 1:1000), anti-PFKP (13389-1-AP, Proteintech, 1:10000), or anti-FLAG (MA1-91878, Invitrogen, 1:10000). Following 1 h of incubation with secondary antibody at room temperature, protein bands were detected using ECL Ultra.

### 5.9. PFKP Enzyme Activity Assay

PFKP enzymatic activity was measured using a commercial PFKP activity assay kit (BC0530, Solarbio) according to the manufacturer’s protocol. Briefly, cell lysates were prepared, and the conversion of fructose-6-phosphate to fructose-1,6-bisphosphate was monitored spectrophotometrically at 340 nm.

### 5.10. Functional Assays

Proliferation was measured with the CCK-8 kit (GK10001-1, GLPBIO) in accordance with the manufacturer’s protocol. At 0, 24, 48, and 72 hours post-treatment (with sodium lactate, sodium oxamate, or control) or post-transfection (with wild-type PFKP, K688R mutant PFKP, or empty-vector plasmids), the absorbance at 450 nm was measured using a microplate reader. We assessed cell migration via a wound-healing assay, recording wound closure at 0, 12, and 24 h. To assess apoptosis, flow cytometry was performed after the cells had been stained using the Annexin V-FITC/PI Apoptosis Detection Kit (G1511-50T, Servicebio).

### 5.11. Statistical Analysis

Data are shown as mean ± SD. A two-tailed t-test was applied for two-group comparisons, and one-way ANOVA was chosen for multiple-group analyses. When two independent variables were present, we employed two-way ANOVA with Tukey’s multiple comparisons test. Spearman’s correlation analysis evaluated the associations between variables. GraphPad Prism 9.5 software was used to perform all statistical calculations. *p < 0.05, **p < 0.01, ***p < 0.001, and ****p < 0.0001 were considered statistically significant.

## 6. Data availability

The datasets in this study are available in the following databases:

6.1. Transcriptomic data: Gene Expression Omnibus GSE21653 (https://www.ncbi.nlm.nih.gov/query/acc.cgi?acc=GSE21653)

6.2. Transcriptomic data: Gene Expression Omnibus GSE25065 (https://www.ncbi.nlm.nih.gov/query/acc.cgi?acc=GSE25065)

6.3. Gene set data: Molecular Signatures Database (MSigDB) (https://www.gsea-msigdb.org/gseasigdb/)

6.4. Lactylation-related gene list: Supplementary Data 1 (included in the supplementary materials of this paper)

## 7. Author contributions

Saifeng Wang: Conceptualization; Methodology; Investigation; Validation; Visualization; Writing—original draft. Zicheng Zhang: Methodology; Project administration; Writing—review and editing. Jiale Chen: Visualization. Minfeng Liu: Supervision; Funding acquisition; Writing—review and editing.

## Contributor Information

Saifeng Wang, Zicheng Zhang, Jiale Chen, Minfeng Liu

## 9. Acknowledgements

This work was supported by the National Natural Science Foundation of China (Grant number: 82274348). We sincerely thank Yukun Cui from Shantou University Medical College, China, for generously providing the HCC1937 cell line.

